# Inflammasome targeting with an NLRP3 agonist therapy is feasible but ineffective in murine hepatocellular carcinoma models with liver damage

**DOI:** 10.1101/2022.06.23.497362

**Authors:** Zhangya Pu, Zelong Liu, Jiang Chen, Koetsu Inoue, Zhiping Ruan, Lingling Zhu, Hiroto Kikuchi, Aya Matsui, Peigen Huang, Dan G. Duda

## Abstract

Co-inhibition of programmed cell death receptor-1 (PD-1) and vascular endothelial growth factor receptor (VEGFR) pathway has shown efficacy in hepatocellular carcinoma (HCC). NLRP3 is a component of the inflammasome involved in the initiation, development, and progression of multiple cancers. We examined whether adding an NLRP3 agonist to dual PD-1/VEGFR inhibitors is feasible and can address treatment resistance in orthotopic HCC in mice with underlying liver damage. Mice with established tumors were treated with an NLRP3 agonist alone, combination of anti-VEGFR2 or the multikinase inhibitor regorafenib with anti-PD1 antibodies, or their combination. In all models tested, NLRP3 agonist therapy showed acceptable toxicity but no effect on tumor growth delay, disease morbidity, or survival. Pharmacodynamic analyses confirmed the effects of NLRP3 agonist therapy on inflammasome, evidenced by a significant elevation in plasma levels of pro-inflammatory cytokines such as IL-1β. However, these changes were not detectable in tumor tissues, where we detected increased expression of immunosuppressive markers IL-6, KC/GRO, CCL9, and IL-18, and immune checkpoint molecules (PD1, PD-L1, and CTLA-4) after NLRP3 agonist therapy. Thus, modulation of the inflammasome with a novel NLRP3 agonist was feasible in mice with orthotopic HCC and liver damage but did not enhance efficacy when combined with anti-PD1/VEGFR therapies.

## Introduction

Hepatocellular carcinoma (HCC) is the most prevalent malignancy of the liver. It is currently ranked a leading cause of cancer-related mortality worldwide and over 850,000 patients are diagnosed with HCC every year [1, 2]. HCC is an aggressive disease occurring predominantly in patients with chronic liver damage caused by viral infections (hepatitis B and C virus), excessive alcohol intake, exposure to aflatoxin, non-alcoholic steatohepatitis (NASH), or metabolic disorders. These factors result in a hepatic environment dominated by inflammation, which increases the risk of carcinogenesis and facilitates the progression of HCCs [3, 4].

In the context of HCC, the presence of a significant immune cell infiltration (referred to as the “immunologically hot” tumor) is generally correlated with a better prognosis, attributed to a greater degree of pre-existing anti-tumor immunity [5, 6]. However, the tumor microenvironment of HCC is usually characterized by immunosuppression, mediated by multiple cellular and biochemical factors involved in immune evasion mechanisms during HCC progression [7, 8]. These mechanisms include the up-regulation of immune checkpoint molecules on HCC and stromal cells promoting the exhaustion of effector immune cells (CD4^+^ and CD8^+^ T lymphocytes), increased infiltration of immunosuppressive cells [tumor-associated macrophages (TAM), T regulatory cells (Tregs), and myeloid-derived suppressor cells (MDSCs)], increased expression of inflammatory cytokines such as IL-6; and the dysfunction of the antigen-presenting process [9-12].

Development of immunotherapy approaches to activate anti-tumor immunity in HCC has seen substantial progress in the last few years with the advent of immune checkpoint blockade (ICB) targeting the programmed cell death receptor 1 (PD-1)/PD ligand 1 (PD-L1) pathway. Monotherapy with anti-PD-1 antibodies (nivolumab, pembrolizumab) was approved by the US

FDA based on phase I/II trials in patients with advanced HCC because of the promising objective response rate but especially due to durable disease control and favorable safety profiles [13] [14]. This pattern of response contrasted to that seen after antiangiogenic therapies, which increased overall survival (OS) but whose benefits were more transient in advanced HCC [15, 16]. Randomized phase III trials of nivolumab and pembrolizumab monotherapy did not reach the prespecified endpoints of survival benefit [17, 18], but a more recent phase III trial of pembrolizumab met the primary endpoint of OS in Asian patients with advanced HCC previously treated with sorafenib (News release. Merck. September 27, 2021. Accessed September 28, 2021).

To enhance the limited efficacy of anti-PD-1 therapy alone, several combinatorial strategies have been developed, including ICB combinations or ICBs with antiangiogenic drugs [8, 19]. We have previously shown that anti-PD-1 antibodies with anti-VEGFR therapies (antibodies or kinase inhibitors) can be effective in murine HCC models, in part by normalizing the abnormal tumor vessels, decreasing the infiltration of Tregs and MDSCs, and enhancing the infiltration and activation of effector CD8 T cells [20] [21]. Recently, a phase III clinical trial (IMbrave150) of combined atezolizumab (an anti-PD-L1 antibody) and bevacizumab (an anti-VEGF antibody) was the first regimen to prolong OS compared to standard antiangiogenic tyrosine kinase treatment with sorafenib [22]. Multiple combinatorial approaches are currently ongoing aiming to establish the efficacy of antiangiogenic tyrosine kinase drugs with anti-PD-1/PD-L1 treatment [19, 23] Yet, despite these significant and exciting new developments, the majority of HCC patients show resistance to these ICB-based approaches, and addressing this unmet need will require new approaches to combat immunosuppression [23].

Inflammation due to liver damage modulates the initiation, development, and progression of carcinogenesis, and involves innate and adaptive immune responses mediated by infiltrating cells such as MDSCs, tumor-associated macrophages (TAMs), or lymphocyte subsets [8, 24, 25]. The inflammatory reaction is mediated by specific cytoplasmic multimetric protein complexes called inflammasomes [26]. Inflammasomes include three key domains of the nucleotide-binding domain (NBD), oligomerization domain (NOD)-like receptors (NLRs), and absent in melanoma 2 (AIM2) and belong to a large family of pattern recognition receptors (PRRs). The PRRs are associated with recognition of pathogen-/ danger-associated molecular patterns (PAMPs and DAMPs) and lead to the activation, maturation, and up-regulation of pro-inflammatory cytokines of IL-1β and IL-18 [27].

Among inflammasome components, the nod-like receptor protein 3 (NLRP3) plays important role in several types of autoimmune and inflammatory diseases such as cold-induced autoinflammatory syndrome (CAPS). Moreover, several recent studies reported that mutation or copy number alteration of the NLRP3 gene was associated with oncogene activation [26, 27]. The function of NLRP3 inflammasome in tumor progression or anti-tumor immunity remains unclear. Constitutive activation of the NLRP3 inflammasome was correlated to the malignant phenotype of human melanoma, lung cancer, and colon carcinoma [27, 28]. Conversely, Wei et al. reported that the expression of all NLRP3 components is either lost or downregulated in the tumor tissues than in the corresponding adjacent non-tumor tissues in HCC [29]. In addition, targeting NLRP3 inflammasome using pharmacological agents may hinder the proliferative and metastatic ability of HCC [29, 30]. Moreover, the specific role of NLRP3 targeting in ICB and antiangiogenic treatment-resistance in HCC is unknown. Here, we examined the feasibility and efficacy of a new NLRP3 agonist alone or with dual anti-PD-1/VEGFR agents in murine HCC models with underlying liver damage.

## Materials and Methods

### Cells and culture condition

Two murine HCC lines were used in the current study: HCA-1 cells, established in the Steele Laboratories on the C3H mouse background [31, 32], and RIL-175 cells, a *p53*-null/*Hras*-mutant line syngeneic to C57Bl/6 mouse background (a kind gift from Dr. Tim Greten, NCI, Bethesda, USA). Cells were cultured in Dulbecco’s Modified Eagle’s Medium (DMEM) (ThermoFisher, USA) was supplemented with 10% fetal bovine serum (FBS) (Hyclone, SH30071.03) and penicillin-streptomycin at a concentration of 100U/ml and 100μg/ml, respectively, at 37°C with 5% CO2. Cells were routinely examined for mycoplasma contamination and authenticated before *in vivo* experiments.

### Orthotopic HCC model

The orthotopic HCC model with liver damage is described in detail elsewhere [33]. Briefly, to induce chronic liver damage, mice were treated three times a week with carbon tetrachloride (CCl4) (Sigma Aldrich, Saint Louis, MO) via oral gavage for 8-12 weeks. One million murine HCC cells in Matrigel (Mediatech/Corning, Manassas, VA) in 1:1 solution were implanted in syngeneic mice (HCA-1 cells in C3H mice and RIL-175 in C57Bl/6). Tumor initiation and growth were monitored using high-frequency ultrasonography twice a week. Mice with established HCC were randomized to the treatment groups when the diameter of the tumor reached approximately 5mm. Per protocol, the experimental endpoint for efficacy studies was moribund status, defined as the following symptoms: significant distress, weight loss of more than 15% compared to pretreatment, body status score >2, and diameter of primary tumor of more 15 mm. All animal experiments were conducted using a protocol approved by the Institutional Animal Care and Use Committee of the Massachusetts General Hospital, Boston, MA.

### Reagents and Treatments

The NLRP3 agonist (BMT-392959) and anti-mouse PD-1 antibody (isotype-matched IgG1:D265A) were all provided by Bristol Myers Squibb Company (USA). These agents were administered as per the manufacturer’s recommendations: NLRP3 agonist was given by intraperitoneal injection (i.p.) at a dose of 10mg/kg once a week, and anti-mouse PD-1 antibody or IgG1 control was administrated at a dose of 10mg/kg via i.p. injections every 4 days. Anti-mouse VEGFR2 antibody (DC101) was purchased from BioXcell (Lebanon, USA) and was given i.p. at a dose of 20 mg/kg three times a week, as described [20]. The multitargeted kinase inhibitor regorafenib was purchased from Selleck Chemicals and administrated at a dose of 10 mg/kg [dissolved in 34% 1,2-propandiol and 34% PEG400 (SigmaAldrich), 12 % pluronic F68 (Thermo Fischer, MA, USA), and 20% purified water] by daily oral gavage, as described [21].

### Proteomic and transcriptomic analyses in time-matched studies in vivo

Blood and tumor samples were collected in separate time-matched experiments using the orthotopic HCC model in C57Bl/6 mice. Tissue collection was performed after 4-hr and 24-hr after treatment with one dose of NLRP3 agonist (10mg/kg) or control in mice with established orthotopic murine RIL-175 HCCs.

To study protein levels of cytokines and chemokines, we separated plasma from the blood samples and extracted proteins from the tissue samples. Briefly, whole blood samples were collected in anticoagulant (EDTA)-coated tubes and processed for plasma separation by centrifugation at 4°C. For protein extraction, each tumor tissue sample was placed in 500 μl/sample lysis buffer. The samples were ground, and then centrifugated at 4°C to collect the liquid phase for protein concentration measurements.

Measurements were performed in duplicate for plasma samples and triplicate for tumor tissue samples using multiplexed array kits from Meso Scale Discovery (Gaithersburg, MD, USA): proinflammatory panel I mouse kit (catalog: K15048D), which includes interferon (IFN)-γ, interleukin (IL)-1β, IL-2, IL-4, IL-5, IL-6, IL-10, IL12p70, TNF-α, and KC/GRO, and Cytokine panel I mouse kit (catalog: K15245D), which includes CCL2, CCL3, CXCL-2, CXCL-10, IL-27p28, IL-9, and IL-33. The procedure was performed as per the manufacturer’s protocols, and plates were analyzed using electrochemiluminescence-based detection on an SQ120 machine.

In addition, total RNA was extracted from the tumor tissue samples using an RNeasy Mini Kit (Qiagen Inc.), and the quality and concentration were analyzed using a NanoDrop Spectrophotometer. Complementary DNA was synthesized using the reverse transcription kit (PrimeScript RT reagent Kit) and quantitative (q)PCR was performed using iTaq Universal SYBR Green Supermix (Bio-Rad Laboratories, Hercules, CA). The housekeeping gene GAPDH was used for reference. The primers used in this study for qPCR are listed in **Table S1**. The relative amplified amount of mRNA was calculated by the 2^−ΔΔCT^ method.

### Statistical analysis

Continuous variables were compared using Student’s t-test with one-tail. When the experiments included more than two groups, the one-way ANOVA with a Brown-Forsythe test was used for multiple comparisons. Categorical variables were analyzed using χ2 (Chi-squared) or Fisher’s test. Kaplan-Meier (K-M) method with Log-rank test was performed to estimate survival probability and the Cox proportional hazard model with a hazard ratio (HR) and 95% CI was executed for statistical survival analysis. All analyses were carried out using GraphPad Prism (version 8.0). A difference was considered statistically significant when P-value was less than 0.05. All studies were conducted at least in triplicate unless otherwise specified.

## Results

### NLRP3 agonist therapy is feasible but is ineffective alone and does not enhance the efficacy of dual VEGFR2 and PD-1 blockade in orthotopic HCC models in mice with liver damage

We first tested the feasibility and efficacy of NLRP3 alone or with dual VEGFR2 and PD-1 blockade in orthotopically grafted RIL-175 murine HCC in C57Bl/6 mice with underlying liver damage. Mice with established tumors (approximately 5mm in diameter) were randomized to one of the four treatment groups: **1**) NLRP3 agonist alone, **2**) anti-VEGFR2 (DC101) and anti-PD-1(aPD-1) antibodies, **3**) NLRP3 agonist combined with anti-VEGFR2 and anti-PD-1 antibodies, or **4**) isotype-matched IgG as a control. All treatments were administered for up to 5 weeks or until mice became moribund (**Fig. 1A**). NLRP3 agonist-related toxicity was evidenced by transient weight loss in the treated groups; the weight loss was less than the 15% limiting toxicity per the animal protocol and all mice recovered 2 days after agent administration (**Fig. S1A**).

**Fig 1.**
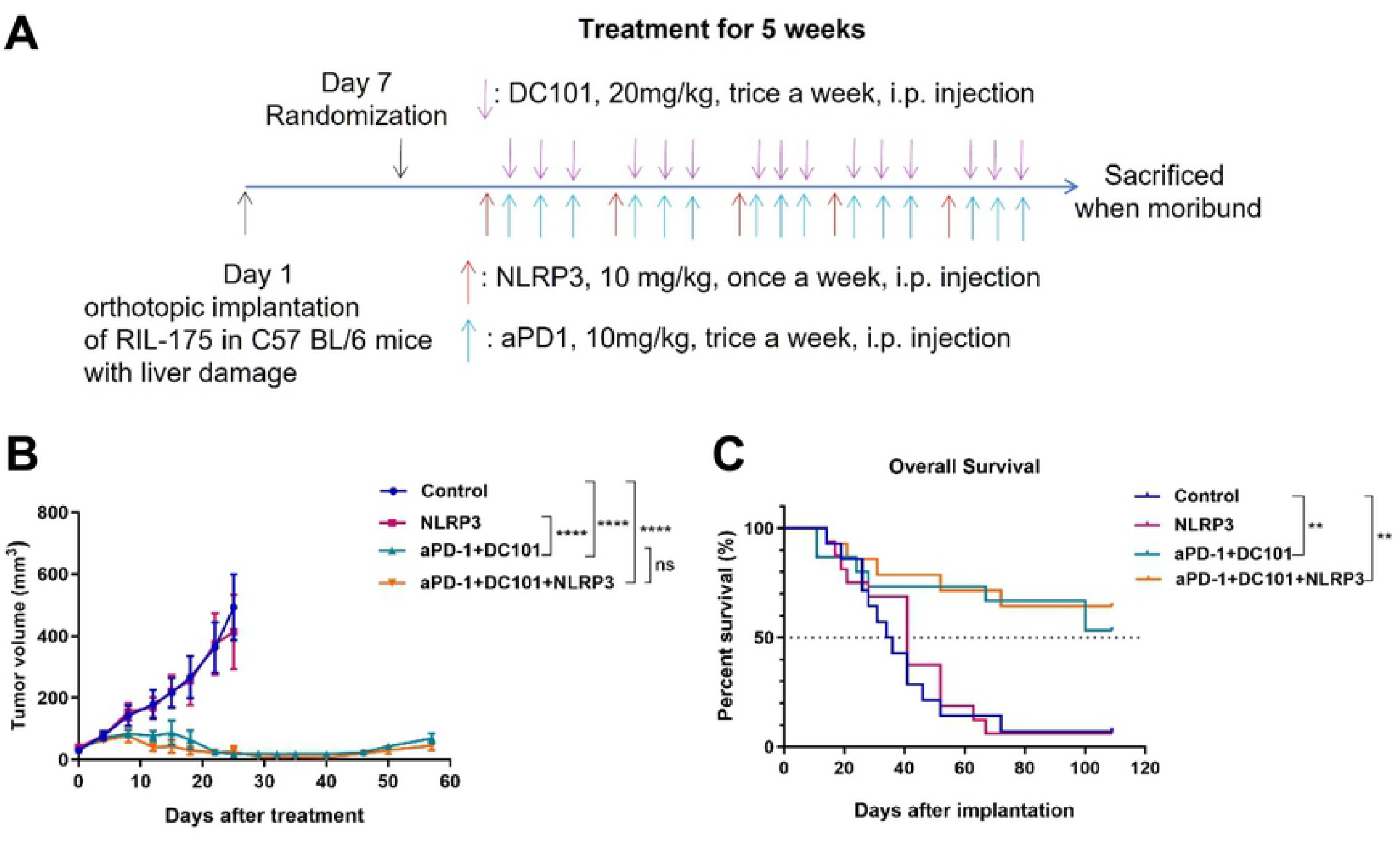
Therapeutic efficacy of NLRP3 agonist alone or combined with anti-PD-1 (aPD-1) and anti-VEGFR2 (DC101) antibodies in orthotopic murine HCC in C57Bl/6 mice. (**A**) Study design. Before intrahepatic implantation of RIL-175 cells, C57Bl/6 mice were treated with CCl4 by gavage for 12 weeks to induce liver damage. (**B, C**) Tumor growth kinetics (**B**) and overall survival distributions (**C**) in the four treatment groups (n=15 mice/group). P values from one-way ANOVA with a Brown-Forsythe test (**B**) and log-rank test (**C**). ** P<0.01, **** P<0.0001, ns: not significant.

As expected, dual antibody blockade of VEGFR2 and PD-1 induced a significant tumor growth delay in this model; in contrast, NLRP3 agonist alone was ineffective, and when combined with dual DC101/anti-PD-1 antibody therapy did not affect tumor growth (**Figs. 1B** & **S1B**). Similarly, we found a significant increase in median OS in tumor-bearing mice after DC101/anti-PD-1 antibody therapy, but no significant difference between NLRP3 alone and control or triple combination versus dual DC101/aPD-1 blockade alone (**Fig. 1C**). In addition to primary tumor growth, mortality in this model can often be caused by the occurrence of ascites and pleural effusions, and less frequently by peritoneal dissemination and lung metastases. Treatment with NLRP3 agonist had no effect on these morbidities and did not impact the effects of dual VEGFR2/PD-1 blockade on these parameters when added to the triple combination regimen (**Fig. S1C-F**).

We next tested the feasibility and therapeutic efficacy of the NLRP3 agonist alone or combined with dual VEGFR2 and PD-1 blockade in the HCA1 murine HCC model in C3H mice with liver damage, which is resistant to anti-PD-1 therapy and highly metastatic to the lungs. As seen in the RIL-175 model, treatment with NLRP3 agonist had acceptable toxicity but showed no benefit when used alone or in combination with dual VEGFR2/PD-1 blockade (data not sown).

Thus, treatment with an NLRP3 agonist was feasible in mouse models of HCC with liver damage, including in combination with VEGFR2 and anti-PD-1 antibodies, but did not confer any additional benefits in reducing mortality or morbidity.

### NLRP3 agonist therapy is feasible but is ineffective alone and does not enhance the efficacy of regorafenib and PD-1 blockade in orthotopic HCC models in mice with liver damage

We first tested the feasibility and efficacy of the NLRP3 alone or with another effective combination approach, using an intermediate dose of the multikinase inhibitor regorafenib (10mg/kg q.d.) and PD-1 blockade in orthotopically grafted RIL-175 murine HCC models in mice with underlying liver damage. Mice with established tumors (approximately 5mm in diameter) were randomized to one of the four treatment groups: **1**) NLRP3 agonist alone, **2**) regorafenib and anti-PD-1 antibody, **3**) NLRP3 agonist combined with regorafenib and anti-PD-1 antibody, or **4**) isotype-matched IgG as a control. All treatments were administered for 3 weeks or until mice became moribund (**Fig. 2A**). NLRP3 agonist and combination therapy with regorafenib/anti-PD-1 antibody showed acceptable toxicity (weight loss less than 15%, and all mice recovered 2 days after agent administration) (**Fig. S2A**).

**Fig 2.**
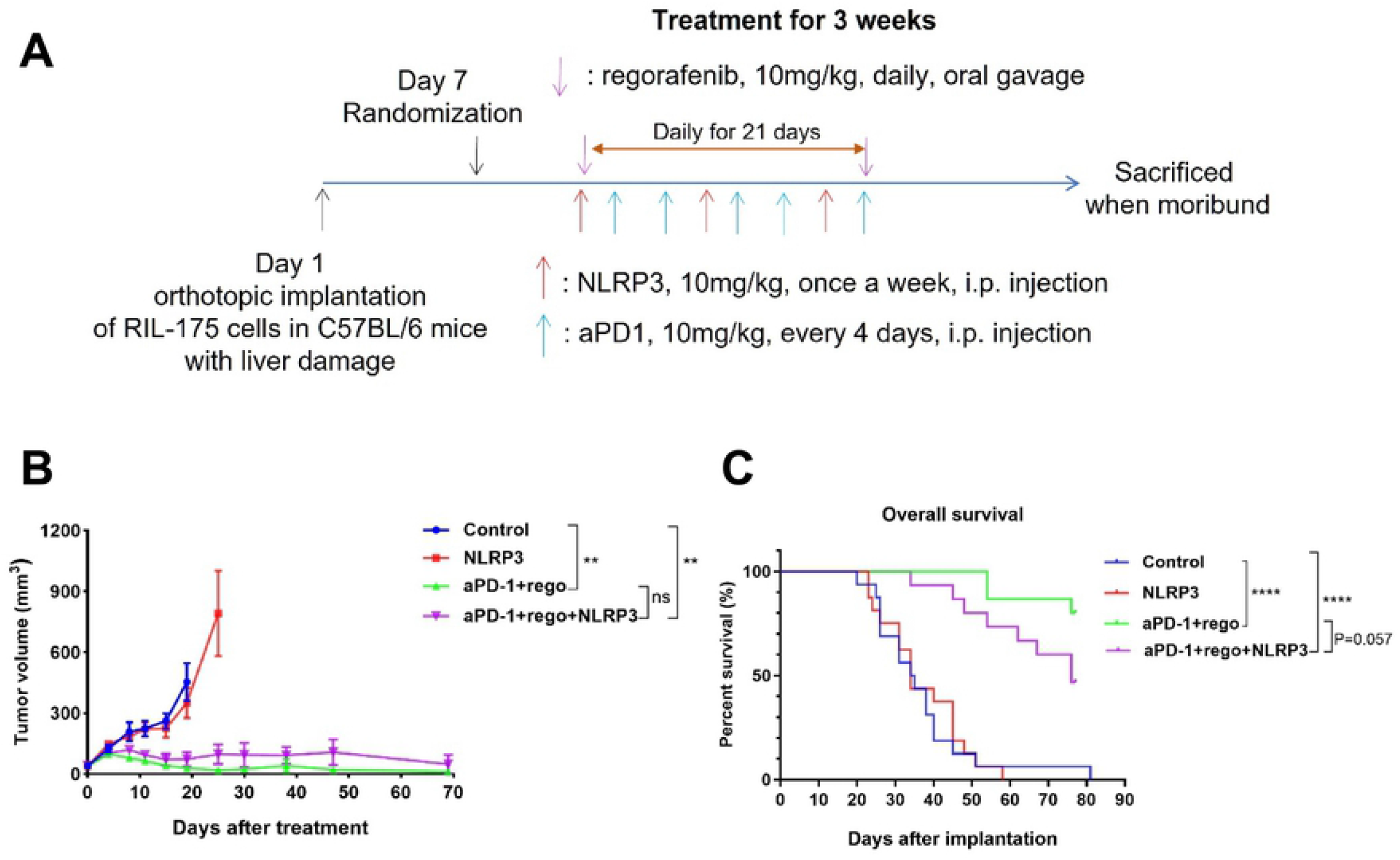
Therapeutic efficacy of NLRP3 agonist alone or combined with anti-PD-1 (aPD-1) and regorafenib (rego) in orthotopic murine HCC in C57Bl/6 mice. (**A**) Study design. Before intrahepatic implantation of RIL-175 cells, C57Bl/6 mice were treated with CCl4 by gavage for 12 weeks to induce liver damage. (**B, C**) Tumor growth kinetics (**B**) and overall survival distributions (**C**) in the four treatment groups (n=15 mice/group). P values from one-way ANOVA with a Brown-Forsythe test (**B**) and log-rank test (**C**). ** P<0.01, **** P<0.0001, ns: not significant.

As seen with dual antibody blockade of VEGFR2 and PD-1, NLRP3 agonist alone was ineffective, and when combined with regorafenib/anti-PD-1 antibody therapy did not increase tumor growth delay and tended to be inferior in terms of median OS compared to dual combination alone (**Figs. 2B-C** & **S2B**). In addition, treatment with NLRP3 agonist did not reduce the incidence of ascites, pleural effusions, peritoneal dissemination, or lung metastasis, when used alone or in combination with regorafenib/anti-PD-1 therapy (**Fig. S2C-F**).

Thus, while treatment with an NLRP3 agonist in combination with regorafenib and anti-PD-1 antibody was feasible in mouse models of HCC with liver damage, it did not confer any benefits in reducing mortality or morbidity.

### NLRP3 agonist therapy increased the expression of immune cytokines in blood circulation but not in the tumor tissues

We next set out to determine whether the lack of efficacy of the NLRP3 agonist was related to the poor pharmacodynamic properties of the novel agent or due to unfavorable biological effects in these models. To this end, we treated mice with established orthotopic RIL-175 murine HCCs with NLRP3 agonist and sacrificed the mice after 4-hr and 24-hr to collect serial blood and tumor tissue samples (**Fig. S3A**). Multiplexed protein array for pro-inflammatory cytokines was performed for plasma and protein extracted from tumor samples to detect changes in the expression level of pro-inflammatory cytokines. The results showed that mice treated with NLRP3 agonist had significant and sustained increases in plasma levels of IL-1β, IL-6, and KC/GRO, while circulating levels of IFN-γ, IL-5, IL-10, and TNF-α were increased only after 4-hr but not after 24-hr (**Fig. 3A**). We detected no differences in IL-2 or IL-12p70 at these time points (**Fig. S3B**).

**Fig 3.**
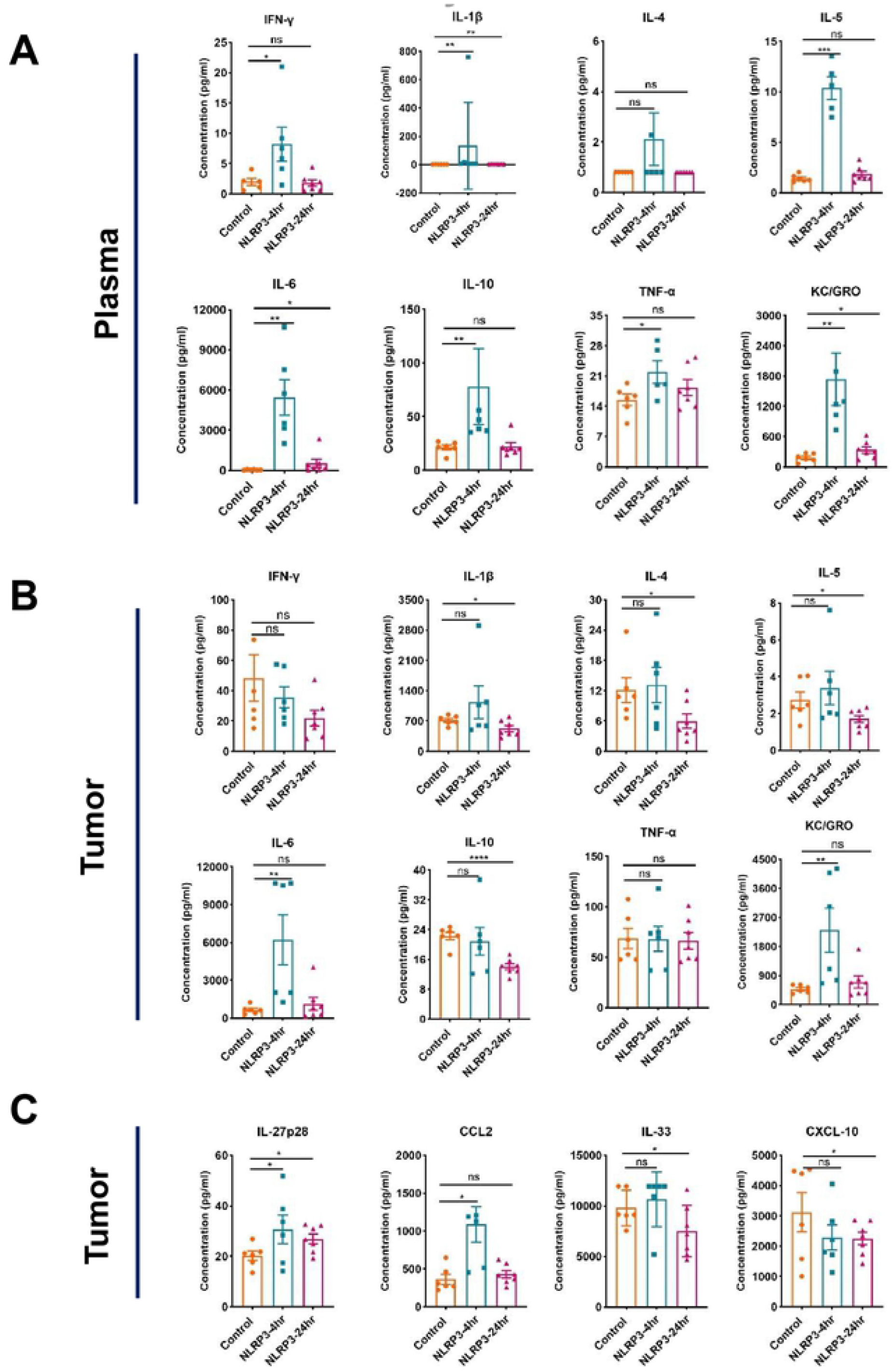
Systemic and intratumoral changes in cytokine/chemokine expression level changed after NLRP3 agonist treatment in mice with orthotopic murine HCC in C57Bl/6 mice. (**A-C**) Multiplexed array measurements of cytokines and chemokines in plasma (**A**) and tumor tissues (**B, C**) at 4 hr and 24 hr after NLRP3 agonist treatment (n=6-7 mice/group). P values from one-way ANOVA with a Brown-Forsythe test. * P<0.05; ** P<0.01, *** P<0.001, **** P<0.0001, ns: not significant.

In tumor tissues, only some of the changes were consistent with the treatment effects detected in blood circulation. Specifically, intratumoral levels of IL-6 and KC/GRO were significantly higher after 4-hr post-NLRP3 agonist treatment compared to control-treated mice. In contrast, intratumoral levels of IL-1β, IL-4, IL-5, and IL-10 were lower after 24-hr post-NLRP3 agonist treatment, and IFN-γ and TNF-α levels were comparable at both time point (**Fig. 3B**). We also used a multiplexed array for chemokine to measure changes in tumor tissues. We found higher levels of IL-27p28 at both time-points and of CCL2 after 4-hr, and decreased IL-33 and CXCL10 levels after 24-hr post-NLRP3 agonist treatment compared to control-treated mice (**Fig. 3C**). There were no differences in the expression levels of CCL-3, IL-9, or CXCL2 in tumors from the NLRP3 treated group at these time points (**Fig. S3C**).

In addition, we used the time-matched RNA extracted from HCC tissue samples to measure changes after NLRP3 treatment in transcriptional levels of selected pro-inflammatory cytokines, TAM and MDSC-related markers, and immune checkpoint molecules using qPCR assay. Among pro-inflammatory cytokines, we found an increased transcriptional level of IFN-γ (both at 4-hr and 24-hr), TGFA (at 4-hr), and CXCL13 (at 24-hr) after treatment with the NLRP3 agonist (**Fig. 4A**). Moreover, the expression levels of CSF1 (at 4-hr), IL-18 and CCL9, and PD-1, PD-L1, and CTLA-4 (both at 4-hr and 24-hr) were up-regulated in samples from NLRP3 treatment group (**Fig. 4B-C**). No changes were detected in the transcriptional levels of IL-1β, IL-5, IL-6, IL-10, IFNA1, CCR2, CXCL10, and CCL26 after NLRP3 agonist treatment at these time points (**Fig. S4**).

**Fig 4.**
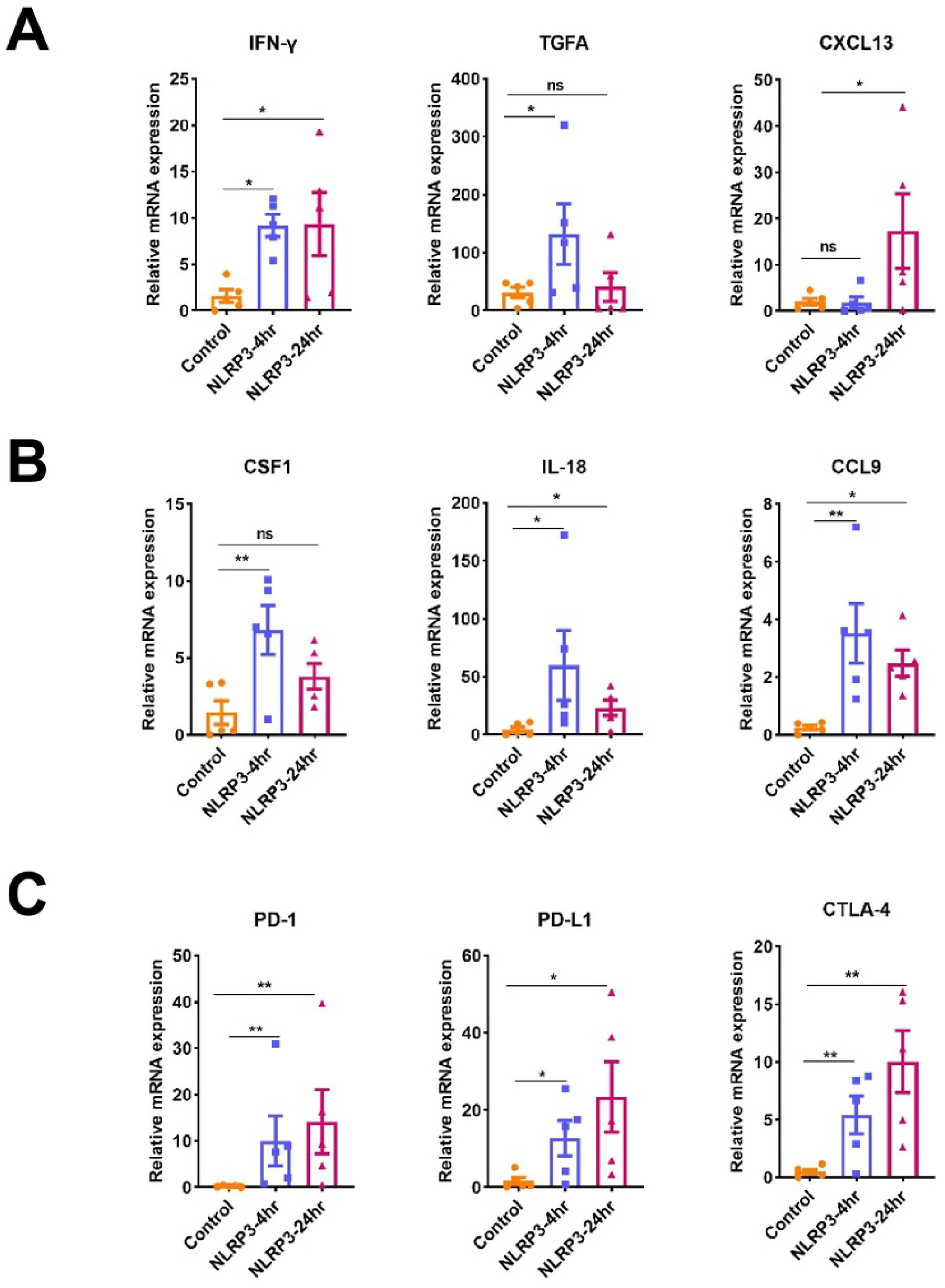
Intratumoral changes in expression levels of selected immunomodulatory genes after NLRP3 agonist treatment in mice with orthotopic murine HCC in C57Bl/6 mice. (**A-C**) Quantitative PCR measurements of proinflammatory factors (IFN-γ, TGFA, CXCL13) (**A**), tumor-associated macrophage (TAM) (CSF1) and myeloid-derived-suppressor cell (MDSC) markers (IL-18, CCL9) (**B**), and immune checkpoint molecules (PD-1, PD-L1, CTLA-4) (**C**), at 4hr and 24 hr after NLRP3 agonist treatment (n=6-7 mice/group). P values from one-way ANOVA with a Brown-Forsythe test. * P<0.05; ** P<0.01, ns: not significant.

## Discussion

New local and systemic treatments have recently become available for advanced HCC patients. However, the increasing incidence of HCC globally and its aggressive progression and refractoriness to treatment make it a persistent health and economic burden. Combination of antiangiogenic drugs and ICB is now a standard modality, and many combinations of kinase inhibitors with anti-PD-1 or anti-PD-L1 antibodies are in advanced stages of clinical development. Based on available data, up to 30% of patients show a response (complete or partial) after these combinational therapies, which in some cases is durable [15, 16, 34, 35]. Thus, despite the impressive efficacy of this combinatorial approach, novel therapeutics are needed to safely extend the benefits of these combinations.

In this study, we tested the feasibility and efficacy of a novel NLRP3 agonist in combination with antiangiogenic agents and anti-PD-1 antibody. Our data confirmed that combining anti-VEGFR2 and anti-PD1 antibodies or regorafenib and anti-PD1 antibodies is effective in murine models, despite using different antibody types, treatment schedules, and suppliers in this study compared to our prior reports [20, 21]. However, although some prior reports supported the approach of targeting NLRP3 to enhance anti-tumor immunity in cancer, our results conclusively demonstrate that this strategy, while feasible, was ineffective in murine HCC models with liver damage [27, 36].

There are several potential explanations for the lack of efficacy for this approach in our models. NLRP3 agonist therapy resulted in high circulating IL-1β and intratumoral IL-18 levels. However, these factors function as a double-edged sword by promoting tumor progression [26]. Indeed, NLRP3 agonist therapy resulted in intratumoral changes at protein level consistent with increased immunosuppression rather than immune activation. While in most contexts these changes did not accelerate progression, there was a tendency for NLRP3 agonist therapy to compromise the efficacy of multikinase inhibitor regorafenib with anti-PD-1 therapy. The immunosuppressive factors included increases in IL-6, IL-10, KC/GRO, IL-27, CCL2, and IL-33 concentrations in HCC tissue. Moreover, the level of CXCL10, a chemokine involved in effector T cell recruitment in this model [21], was reduced by NLRP3 agonist therapy. Of note, expression levels of CSF-1 and CCL9 were increased at the transcriptional level after NLRP3 agonist therapy, along with immune checkpoint molecules (PD-1, PD-L1, and CTLA-4), potentially indicating accumulation or activation of myeloid cells (MDSCs and TAMs). Irrespective of the mechanisms involved, our preclinical results do not support the use of this strategy alone or with antiangiogenic agents and anti-PD-1 antibodies in HCC patients.

Future approaches will need to address two major unmet needs in ICB-based therapy for HCC. One is addressing the profound immunosuppression in the microenvironment of advanced HCC, which likely limits treatment efficacy. Another unmet need is developing approaches to directly target the cancer cells as an approach to enhance the efficacy of ICB with antiangiogenic therapy. To address these problems, multiple new strategies are being currently developed. For example, we have recently shown that judicious scheduling of anti-PD-1 antibody with multikinase inhibitors can show efficacy while reducing drug exposure [21]. Our prior work also demonstrated the favorable reprogramming of the HCC microenvironment when anti-VEGFR and anti-PD-1 therapy was combined with CXCR4 inhibition [37]. Moreover, our group has also recently demonstrated the efficacy of targeting HCC using p53 mRNA therapy to enhance the efficacy of anti-PD-1 therapy [38]. Others have shown the potential therapeutic usefulness of targeting the DNA-activated STING pathway combined with anti-PD-1 therapy in HCC [39]. There is increasing interest in targeting innate or innate-like cells, recently reviewed in HCC [40], or gut microbiome in HCC, as discussed by Schwabe RF et al [41]. Finally, many efforts are directed toward the development of epigenetic modifiers such as HDAC inhibitors as immunomodulators for ICB therapy in HCC models [42], and are currently being tested in clinical trials in other cancers [43].

## Conclusions

NLRP3 agonist showed acceptable toxicity but lacked efficacy in orthotopic murine HCC models in mice with underlying liver damage and did not show any additional benefit when combined with effective anti-PD-1/anti-angiogenic agents. While NLRP3 agonist therapy was associated with pharmacodynamic increases in blood IL-1β levels, the protein and transcriptional analyses revealed an increase in immunosuppressive factors in the HCC tissues. Our preclinical study data do not support the further development of NLRP3 agonists in HCC patients.

## Acknowledgments

The authors would like to thank Mark Duquette, Anna Khachatryan, and Sylvie Roberge for outstanding technical support.

## Author contributions

**Conceptualization**: Dan G. Duda

**Data curation**: Dan G. Duda, Zhangya Pu

**Formal analysis**: Zhangya Pu

**Funding acquisition**: Dan G. Duda

**Investigation**: Zhangya Pu, Zelong Liu, Jiang Chen, Koetsu Inoue, Zhiping Ruan, Lingling Zhu, Hiroto Kikuchi, Aya Matsui

**Project Administration**: Dan G. Duda **Resources**: Peigen Huang **Supervision:** Dan G. Duda **Validation:** Zhangya Pu **Visualization:** Zhangya Pu

**Writing – original draft preparation:** Zhangya Pu

**Writing – review & editing:** all authors

### Funding

This study was supported through a sponsored research agreement with Bristol Meyer Squibb, who provided funds and reagents (to DGD). DGD’s research is supported through NIH grants R01CA254351, R01CA260857, R01CA247441, P01CA261669, and R03CA256764, and

Department of Defense grants PRCRP W81XWH-19-1-0284 and PRCRP W81XWH-21-1-0738.

### Informed Consent Statement

Not applicable.

### Data Availability Statement

The original contributions presented in this study have been already included in the main text or supplementary materials. For any inquiries, please contact the corresponding author.

### Conflict of Interest Disclosure

DGD received consultant fees from Innocoll and research grants from Exelixis and Surface Oncology. No reagents from these companies were used in this study.

## Supporting information

**Fig. S1: Effects of NLRP3 agonist alone or combined with anti-PD-1 (aPD-1) and anti-VEGFR2 (DC101) antibodies in orthotopic murine HCC in C57Bl/6 mice. A** Changes in body of mice during treatment in the overall cohorts and in individual mice in the four treatment groups (n=15 mice/group). **B** The tumor volume curve in individual mice. **C-F** Incidence of ascites **C**, pleural effusion **D**, lung metastasis **E**, and peritoneal dissemination **F** in the in the four treatment groups (n=15 mice/group).

**Fig. S2: Effects of NLRP3 agonist alone or combined with anti-PD-1 (aPD-1) and regorafenib (rego) in orthotopic murine HCC in C57Bl/6 mice. A** Changes in body of mice during treatment in the overall cohorts and in individual mice in the four treatment groups (n=15 mice/group). **B** The tumor volume curve in individual mice. **C-F** Incidence of ascites **C**, pleural effusion **D**, lung metastasis **E**, and peritoneal dissemination **F** in the in the four treatment groups (n=15 mice/group).

**Fig. S3: Systemic and intratumoral changes in cytokine/chemokine expression level after NLRP3 agonist treatment in mice with orthotopic murine HCC in C57Bl/6 mice. A** Experimental design. **B, C** Multiplexed array measurements of cytokines and chemokines in plasma **B** and tumor tissues **(B and C)** at 4hr and 24 hr after NLRP3 agonist treatment. Statistical analysis using one-way ANOVA with a Brown-Forsythe test, ns: not significant.

**Fig. S4: Intratumoral changes in expression levels of immunomodulatory genes after NLRP3 agonist treatment in mice with orthotopic murine HCC in C57Bl/6 mice**. Quantitative PCR measurements of proinflammatory factors (IL-1β, IL-5, IL-6, IL-10, IFNA1), tumor-associated macrophage (TAM) (CCR2, CXCL10) and myeloid-derived-suppressor cells (MDSC) (CCL26) markers at 4hr and 24 hr after NLRP3 agonist treatment (n=6-7 mice/group). Statistical analysis using one-way ANOVA with a Brown-Forsythe test, ns: not significant.

